# Millisecond Prediction of Protein Contact Maps from Amino Acid Sequences

**DOI:** 10.64898/2026.03.15.711852

**Authors:** Runfeng Lin, Sebastian E. Ahnert

## Abstract

Protein structure prediction typically outputs static coordinates, often obscuring the underlying physical principles and conformational flexibility. In this work, we present a coarse-grained generative framework to recover the Circuit Topology (CT) of proteins using Generative Flow Matching. We represent protein architecture using highly compressed Secondary Structure Elements (SSEs), reducing the sequence length to roughly 1/13 of the original amino acid sequence. We show that this minimal representation captures the essential “topological fingerprint” required to determine the global fold. By employing a joint-prediction head, our model simultaneously generates contact probabilities and asymmetric topological features, achieving a mean F1 score of 0.822 at the SSE level. Notably, our results demonstrate a counter-intuitive robustness in capturing long-range interactions, suggesting that global topology acts as a stable constraint compared to local residue packing. Furthermore, we show that these coarse-grained predictions can be mapped back to residue-level contact maps with sub-helical precision, yielding a mean alignment error of 2.69 residues. The probabilistic nature of the flow model effectively separates the stable structural signal of the folding core from flexible regions, providing a physically interpretable view of the protein’s conformational ensemble. This pipeline is extremely fast, capable of completing a contact map prediction from amino acid sequence in an average of 110 milliseconds on a single GPU. These ultra-fast and accurate predictions provide a valuable tool for identifying conserved protein folding cores, facilitating the exploration of the protein structural genotype-phenotype (GP) map through large-scale sampling of mutants with highly similar folding cores.

## 1 Introduction

Proteins serve as the fundamental building block of life, deriving their diverse functions from intricate three-dimensional structures. While local stability is governed by electrostatic forces, van der Waals interactions, and hydrogen bonding,[1] the global fold is determined by long-range constraints. Conventionally, structural similarity is quantified using geometric metrics such as RMSD or TM-align.[2] However, these rigid-body superposition methods often overlook the fundamental topological constraints that govern the folding process. A protein may undergo significant geometric fluctuations while maintaining a stable topological state, or conversely, small geometric changes can lead to distinct topological entanglements.[3]

To address the limitations of geometric descriptors, the framework of Circuit Topology (CT) was developed to provide a rigorous mathematical description of folded linear chains.[3, 4] Unlike knot theory, which primarily applies to closed loops, CT classifies intra-chain contacts in open chains into three fundamental arrangements: Series (S), Parallel (P), and Cross (X).[3] Theoretical studies have demonstrated that these topological relations impose strict constraints on the conformational search space, effectively reducing the entropic cost of folding.[5] Specifically, non-trivial topologies such as Parallel and Cross motifs have been shown to influence folding kinetics through mechanisms like the “zipping effect,” distinguishing physically accessible native folds from random globule states.[1, 5] Furthermore, topological descriptions offer a robust way to categorize folding pathways and intermediate states that are invisible to standard coordinate-based analysis.[6]

These global topological rules are intrinsically linked to the hierarchical arrangement of local Secondary Structure Elements (SSEs). Previous research established that local secondary structures significantly restrict the conformational space,[7, 8, 9, 10] serving as nucleation sites for global topology. Recently, Lin et al.[11, 12] demonstrated that protein structures can be efficiently compressed into SSE sequences (averaging 1/13 of the amino acid sequence length) while retaining critical structural information. This suggests that the pairwise organization of SSEs acts as a “topological fingerprint” of the protein fold. However, proteins are not static entities; they exist as thermodynamic ensembles with intrinsic conformational flexibility.[13] Proteins sharing high sequence identity or SSE similarity may adopt distinct structural states due to changes in relative orientation or loop dynamics. Traditional deterministic models often fail to capture this structural plasticity, collapsing the diverse conformational landscape into a single mean structure. To bridge this gap, we employ a Generative Flow Matching Model,[14] adapted with a BERT-style architecture,[15] to model the conditional probability distribution of SSE arrangements. Unlike deterministic predictors, this generative approach allows us to capture both the inter-SSE relationships and the inherent uncertainty associated with conformational flexibility.

While Circuit Topology offers a robust abstraction superior to atomic coordinates, predicting the full topological matrix directly is computationally prohibitive. A complete topological description involves a matrix of relations with dimensions scales with the fourth power of the number of SSEs (*L*^4^), which is highly sparse, noisy, and computationally expensive to learn. Consequently, rather than predicting the high-dimensional circuit topology directly, we propose a proxy approach: predicting the contact map and the corresponding fractional contact positions. This strategy allows us to derive the underlying circuit topology (CT) efficiently, leveraging the generative model to resolve the ambiguity in flexible regions while maintaining rigorous topological constraints. We validated our method using secondary structures extracted from both experimental structures and Porter 6 [16] predictions. The high accuracy across both inputs suggests that our model learns the underlying topological principles of protein folding from SSEs, allowing it to tolerate small variations in the SSE string rather than overfitting to exact sequence matches. By achieving similar accuracy using secondary structures predicted from amino acid sequences, this framework opens a new avenue for studying the protein structural genotype-phenotype (GP) map[17] through largescale sampling of protein sequences with highly conserved CT-based folding cores. The complete workflow is illustrated in Figure 1.

**Figure 1.**
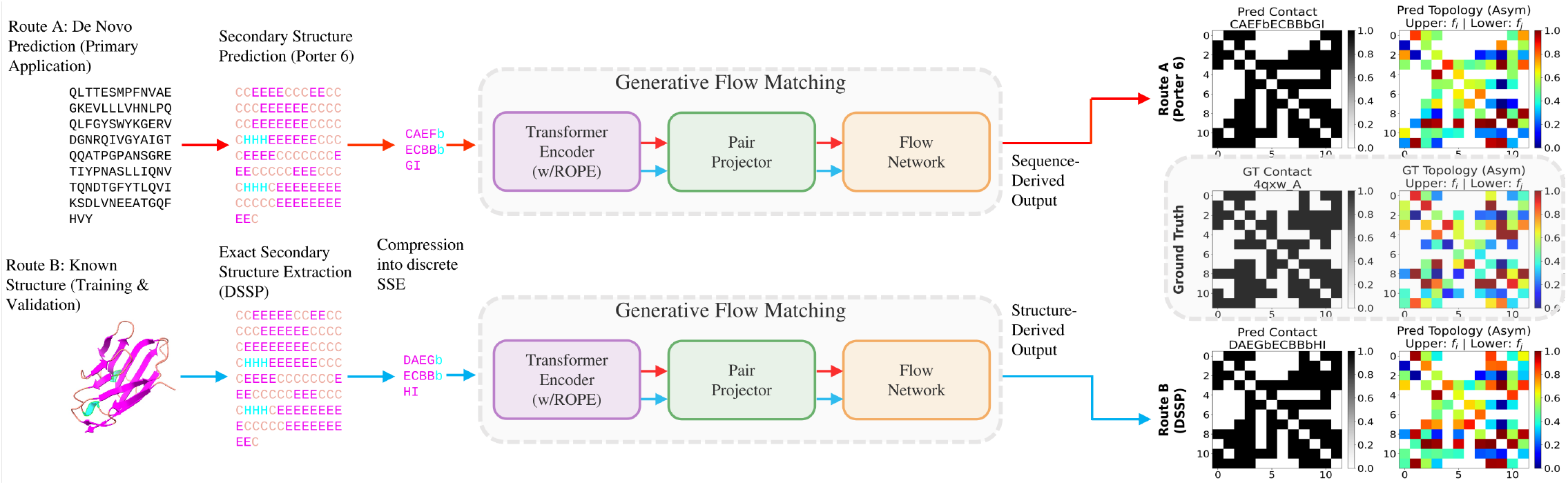
Workflow for CT prediction. **Left:** Secondary structure extraction for protein 4qxw A via two independent pathways: sequence-based 1D prediction using Porter 6 (Route A, orange) and 3D experimental assignment using DSSP (Route B, blue). **Center:** The shared generative flow matching neural network architecture that processes the discrete SSE inputs. **Right:** Comparison of the resulting predicted contact and topology maps from both routes against the experimental Ground Truth (GT).

## 2 Results

### 2.1 Performance of Coarse-Grained Contact Prediction

The model was evaluated on the RCSB dataset, filtered to maintain a maximum of 90% sequence identity with the training set. We first assessed the prediction accuracy using the F1 score at the SSE level. The results demonstrate a high degree of agreement between predictions and ground truth, achieving an average F1 score of 0.822 across the test set (Figure 2). This performance corroborates our previous findings that compact SSE representations provide sufficient constraints to define the global protein fold and structural properties.[11, 12].

**Figure 2.**
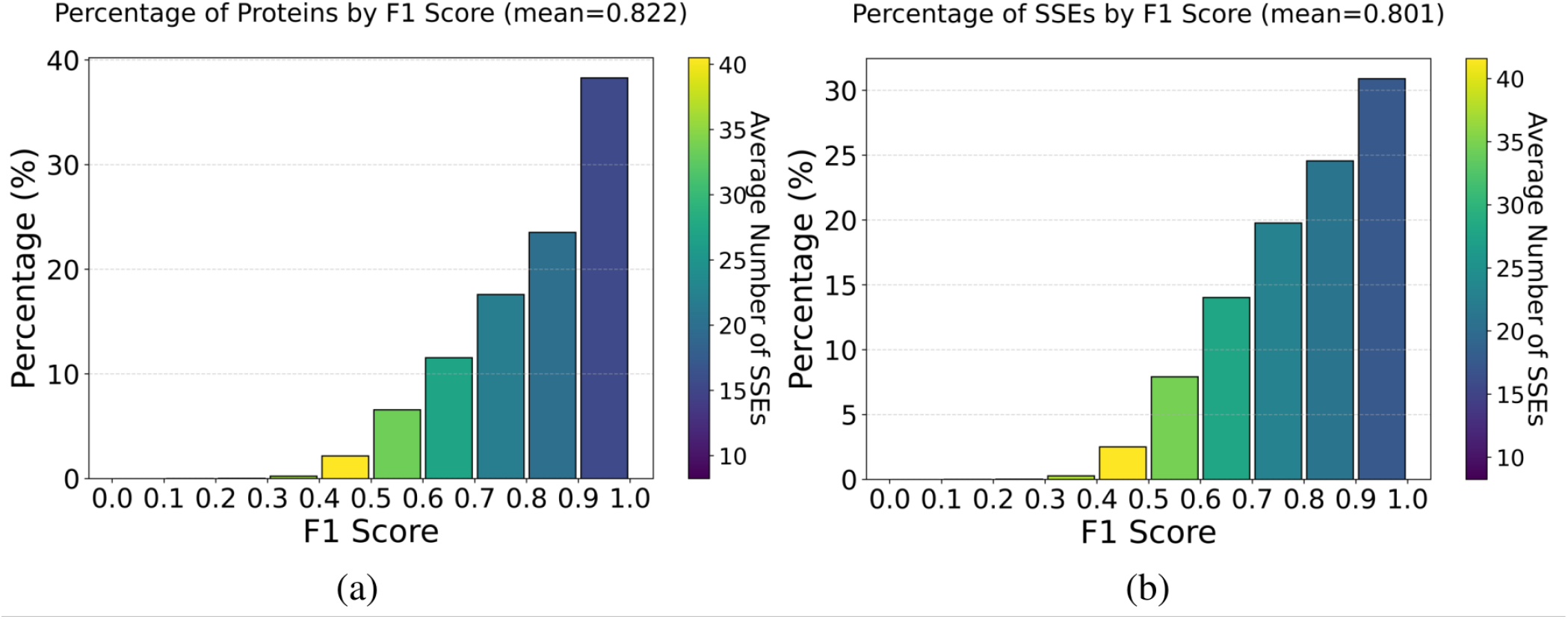
Performance distribution of coarse-grained contact map predictions. **(a)** Histogram of F1 scores for SSE-level contact prediction by proteins (mean = 0.822). **(b)** Histogram of F1 scores for SSE-level contact prediction by SSEs (mean = 0.801). The color bar indicates the average number of SSEs. Smaller proteins (blue, *<* 30 SSEs) generally achieve higher accuracy than larger, complex structures (yellow).Though some proteins have the same SSEs, but the overall F1 score by proteins or by SSEs are almost the same suggest model is not biased toward specific SSEs.

We further analyzed model performance by protein size. As illustrated by the colour gradient in Figure 2, the model performs particularly well on proteins with fewer than 30 SSEs. Given that the coarse-grained SSE representation compresses the sequence length by a factor of approximately 13, a threshold of 30 SSEs effectively encapsulates the typical size range of single and double domains (50–200 residues). Consequently, our method covers the biologically relevant topological space for the majority of protein domains. The observed performance attenuation in larger structures is likely attributable to the increased complexity and conformational flexibility characteristic of multi-domain proteins.

To further probe the model’s capability in capturing global geometry, we examined prediction accuracy across different interaction ranges. Traditional residue-level methods often struggle with long-range interactions due to the restricted receptive fields of CNNs[18] or information loss in RNNs[19]. To evaluate whether our coarse-grained approach overcomes this limitation, we adapted classical residue-level separation criteria [20] to the topological domain. Interactions were classified based on SSE index separation (*k*) into three categories: short-range (*k* ≤ 2), medium-range (3 ≤ *k <* 5), and long-range (*k* ≥ 5).

The results, presented in Figure 3, reveal a counter-intuitive robustness in long-range interactions (Mean F1 = 0.818). Unlike traditional methods where accuracy typically decays with increasing distance, our model performs exceptionally well in the global regime [20, 21]. This indicates that the model has effectively learned the global folding logic and stable hydrophobic core formation, rather than merely memorizing local secondary structure packing (short-range). Furthermore, the relative dip in performance for medium-range interactions likely reflects the intrinsic conformational flexibility of intermediate regions (such as loops or domain boundaries) [13], which are physically less constrained than the folding core.

**Figure 3.**
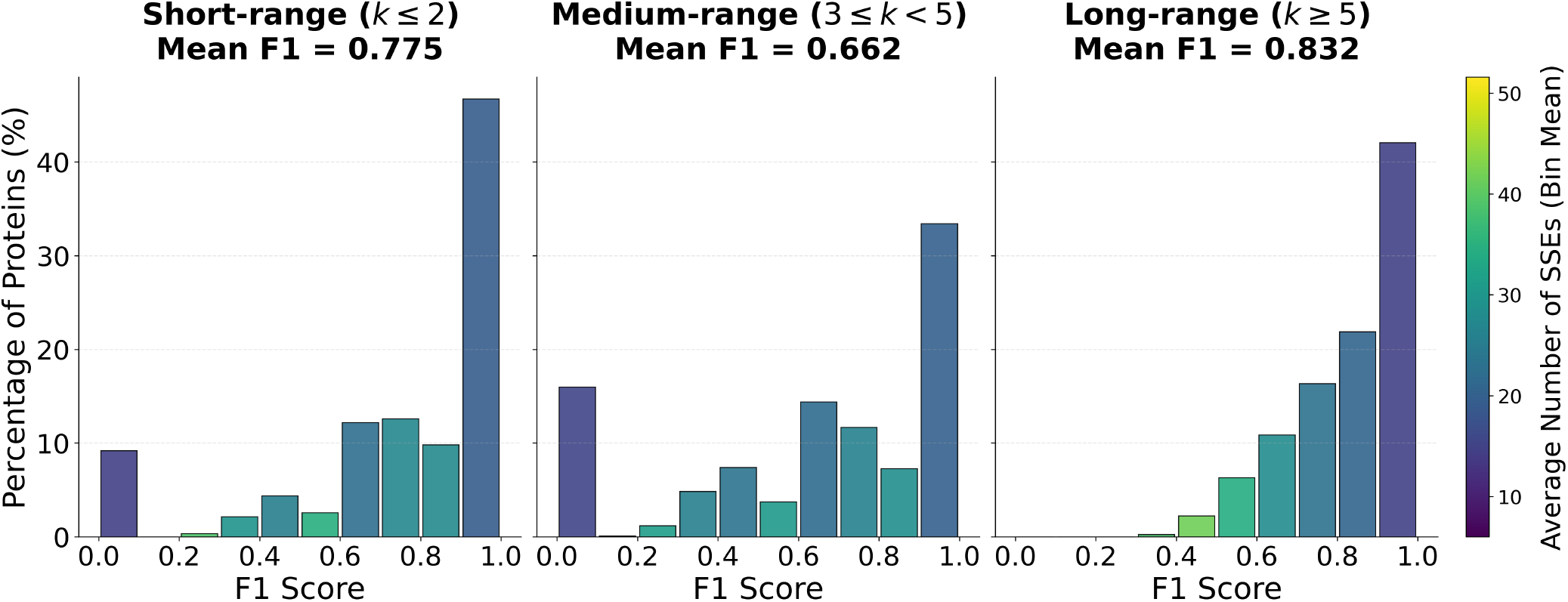
Performance distribution of coarse-grained contact map prediction across different topological ranges. The interactions are stratified by SSE index separation *k* into short-range (*k* ≤ 2), medium-range (3 ≤ *k <* 5), and long-range (*k* ≥ 5). The histograms show the percentage of proteins (y-axis) distributed by F1 score (x-axis). The color scale represents the average protein size (number of SSEs). Notably, the model demonstrates superior robustness in capturing long-range global topology compared to local interactions.

### 2.2 Topological Fidelity and Localization Precision

To validate the robustness of our topology recovery, we evaluated the global connectivity of the protein fold using Circuit Topology (CT), which classifies contact pairs into Series (S), Parallel (P), or Cross (X) arrangements. This representation is invariant to local geometric fluctuations but highly sensitive to global misfolding [1, 3].

We first quantified topological agreement using the Jaccard index. At the SSE level, the model achieved a mean Jaccard index of 0.57. This result reflects the quadratic nature of topological relations, where a single missing contact disrupts all associated topological pairs. To further evaluate the structural “skeleton,” we utilized the Damerau-Levenshtein (DL) similarity to measure the editing distance between topological sequences. The model achieved a Macro-DL similarity (SSE-level) of 0.851 and a Micro-DL similarity (residue-level contact centers) of 0.693. When conditioned strictly on True Positive (TP) contacts, the DL similarity increases to 0.785, indicating that once a contact is correctly identified, its relative topological placement is recovered with high accuracy. A visual representation of this high topological agreement is provided in Figure 5, which illustrates the alignment between ground truth and predicted connectivity for both Macro- and Micro-level circuit topologies on a representative protein.

As shown in Figure 4(a), the model performance follows a physical hierarchy: Series (*S*, recall=0.79) is the most accessible, followed by Parallel (*P*, recall=0.69) and Cross (*X*, recall=0.64). Notably, the high recall for Cross topologies—the most complex entanglements in protein folds suggests that the generative model learns global physical constraints rather than merely memorizing local packing.

**Figure 4.**
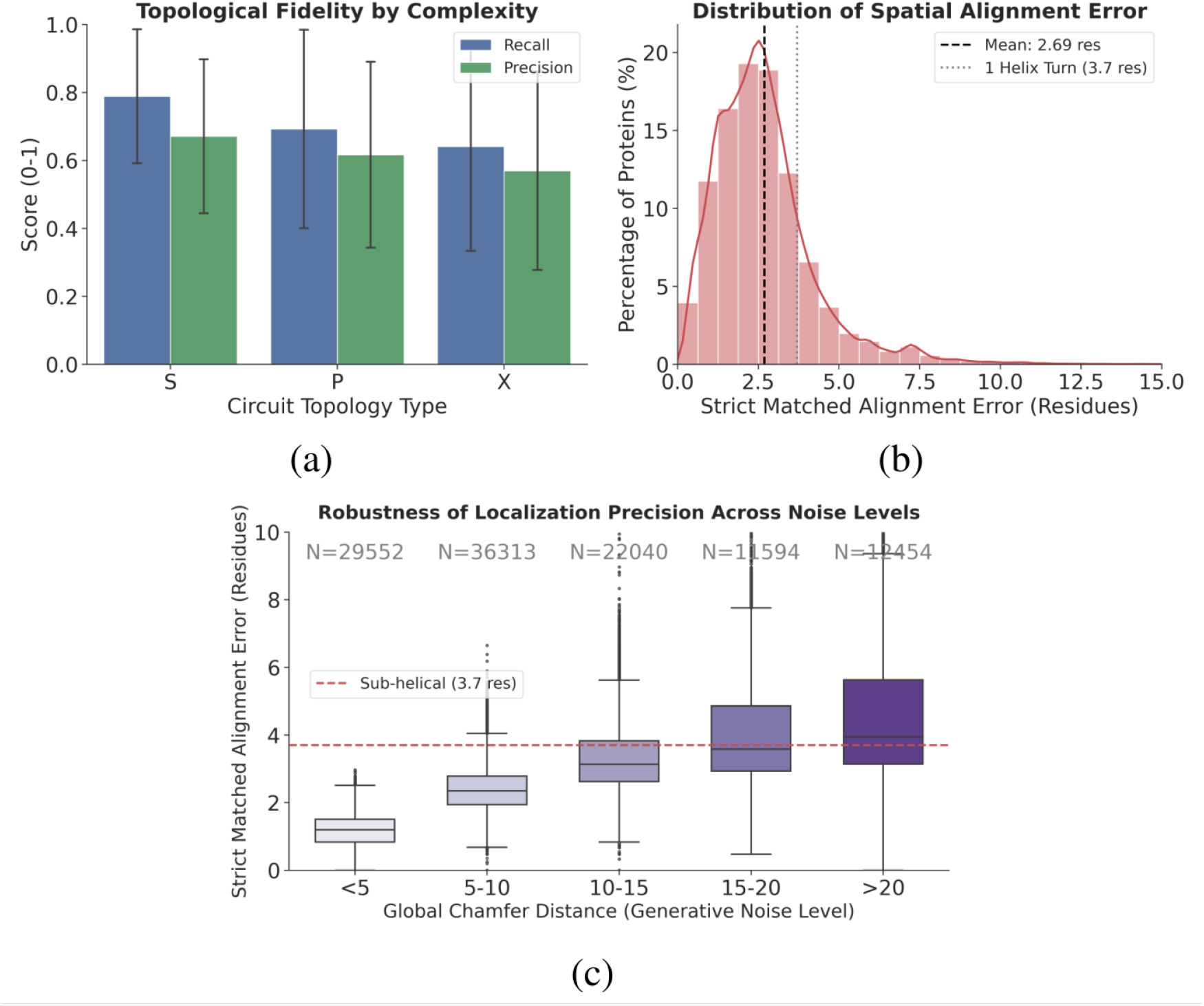
Quantitative Assessment of Topological Robustness. **(a)** Topological fidelity across different circuit arrangements. The model maintains high recall for complex Cross (X) entanglements, indicating successful recovery of global folding logic. **(b)** Distribution of spatial alignment error for matched contacts. The mean error of 2.69 residues falls well within the sub-helical precision threshold (3.7 residues), ensuring the physical viability of the predicted interaction interfaces. **(c)** Robustness of localization precision across noise levels. Boxplots show that even for predictions with high generative noise (Global Chamfer Distance *>* 20), the median localization error of true positive contacts remains robustly below the sub-helical limit.

**Figure 5.**
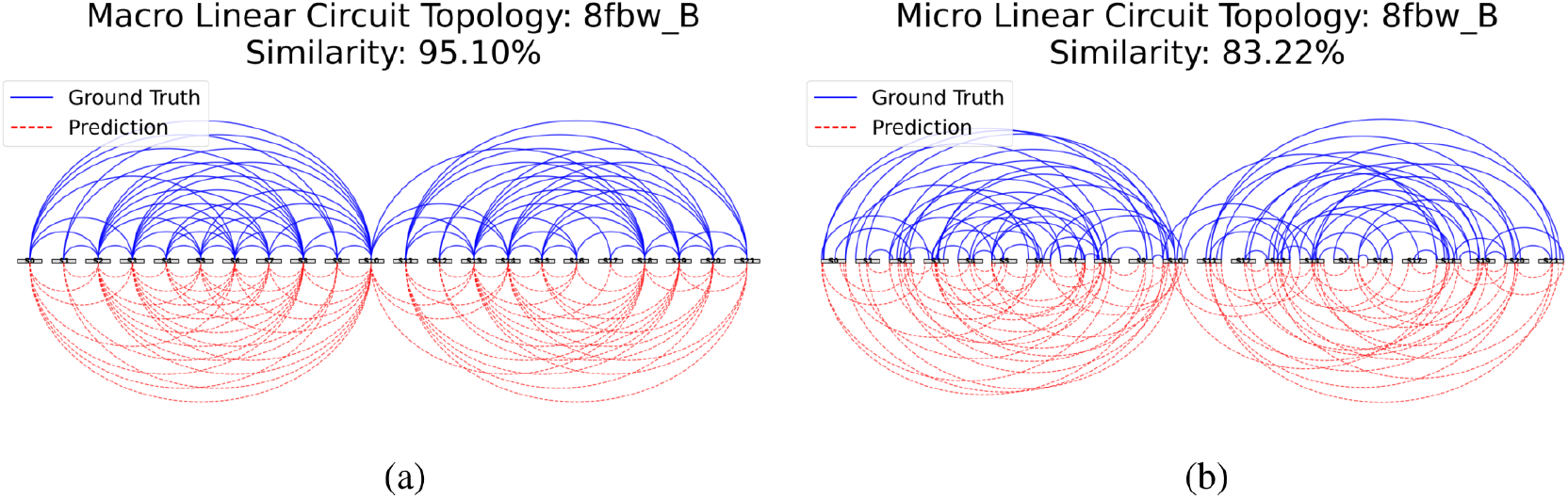
Linear Circuit Topology Visualization for Protein 8fbw B. The diagrams illustrate the alignment between Ground Truth (top, blue solid lines) and Predicted (bottom, red dashed lines) topological connections. **(a)** Macro-Topology: Visualizes aggregated SSE-level connectivity, achieving a topological similarity of 95.10%. **(b)** Micro-Topology: Visualizes residue-level contact center alignments, reflecting the higher-resolution spatial registration with a similarity of 83.22%. The close agreement in arc nesting and crossing patterns demonstrates the model’s ability to capture the hierarchical entanglement of the fold.

A key challenge in mapping coarse-grained predictions to residue-level coordinates is residue displacement. We quantified this using “Strict Matched Alignment Error” (MAE). As shown in Figure 4(b), the mean displacement is 2.69 residues, which is significantly below the 3.7-residue threshold of a single *α*-helical turn [22], implying that identified topological contacts are localized with near-atomic fidelity.

Finally, we assessed the decoupling of generative noise from structural signal. As illustrated in Figure 4(c), while the total background noise (Global Chamfer Distance) varies[23], the localization error for true positive contacts remains stable and predominantly within the sub-helical precision zone (*<* 3.7 res). This suggests the generative flow effectively separates the “signal” of the folding core from the “noise” of uncertain, flexible regions.

While a random null model suggests an equal 33.3% probability for each topological arrangement, sequential constraints in physical chains further reduce the likelihood of complex entanglements. Specifically, “Cross” motifs require the precise interleaving of two long-range interactions, making them statistically much rarer (around 8.9% in the whole RCSB database) than simple local contacts. Despite this low random probability, our model achieves a high recall of 0.64 for Cross (X) topologies. This significant deviation from the random baseline demonstrates that the generative flow effectively captures the non-random, evolutionarily conserved structural logic of the protein fold, rather than merely reflecting accidental proximity.

### 2.3 Residue-Level Reconstruction and Geometric Fidelity

To bridge the gap between coarse-grained topological predictions and residue-scale interactions, we developed a reconstruction protocol designed to prioritize robust structural features over transient noise. Rather than attempting to reconstruct the complete, high-resolution residue contact map—which often contains dispensable weak links—we focused on recovering the essential interaction cores.

On the ground-truth (GT) side, contacts were processed via binary dilation and clustering to merge proximal residues within a 2-pixel radius into coherent patches. This step effectively filters out minor fluctuations and aligns the evaluation with the model’s preference for strong, physically significant connections. On the prediction side, SSE-level constraints were projected into residue-space “seeds” and smoothed using a Gaussian kernel (*σ* = 1.5). This “blob” representation introduces a secondary layer of coarse-graining, allowing for a robust assessment of interaction loci without penalizing minor atomic displacements.

Importantly, when evaluated under this robust protocol, the agreement was even higher than the SSE-level metrics. As shown in Figure 6(a), the residue-level reconstruction achieved a mean F1 score of 0.840. This improvement upon mapping back is noteworthy; it suggests that misclassifications at the SSE level often occur at the “fuzzy boundaries” of secondary structures rather than misidentifying the core interaction topology. By strictly constraining the residue-level map to the predicted SSE pairs, our method effectively filters out background noise, ensuring that the reconstructed contact map retains a high signal-to-noise ratio.

**Figure 6.**
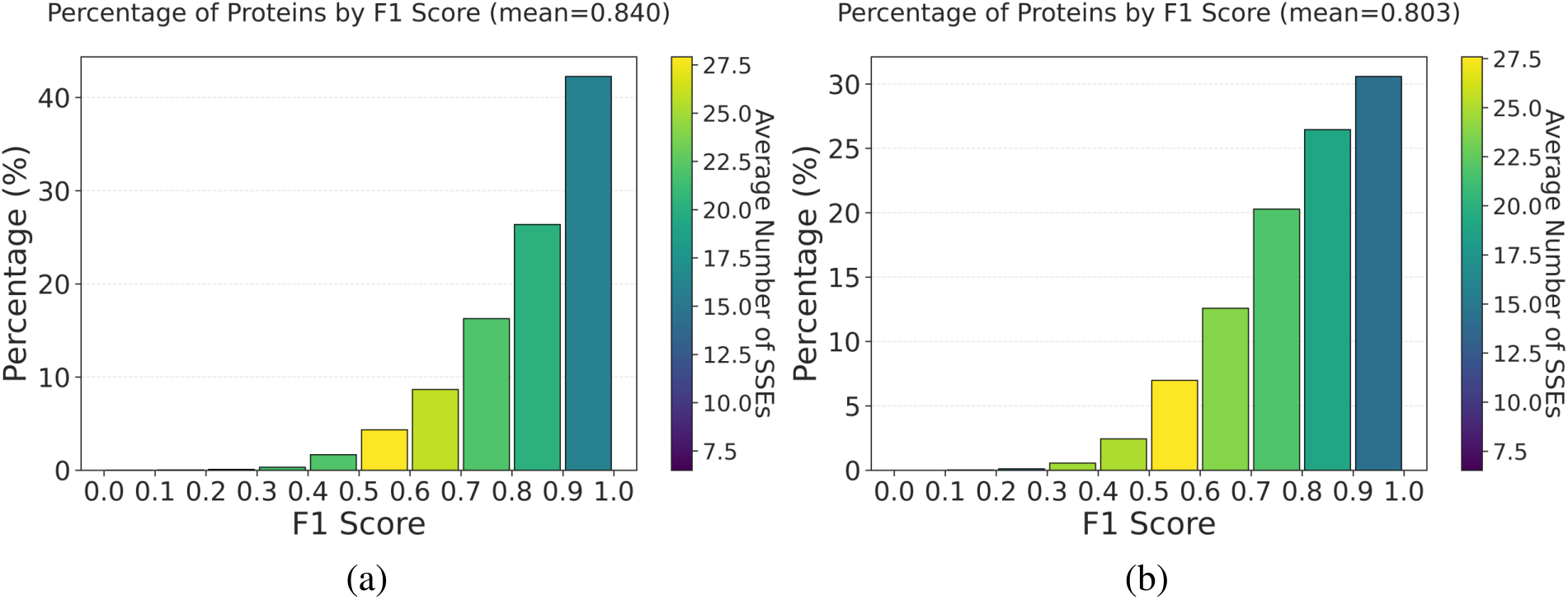
Residue-level reconstruction accuracy. **(a)** Distribution of F1 scores for residuelevel contact maps reconstructed from experimental SSE inputs (mean = 0.840). The higher mean compared to SSE-level prediction suggests the model effectively captures the contact skeleton. **(b)** Distribution of F1 scores for residue-level contact maps reconstructed from Porter 6 predicted SSE inputs (mean = 0.803). Performance is negatively correlated with protein size (indicated by colour).

To further validate the generalization capability of the model, we replaced experimental secondary structures with predictions from Porter 6 [16]. Despite the inherent error propagation from Porter 6 (which achieves ~84–86% accuracy), the degradation in our topological prediction was minimal, with the mean F1 score dropping only by ~0.04 to 0.803 (Figure 6(b)). This demonstrates that our model successfully learns the global folding logic rather than relying on perfect local definitions, making it highly robust for real-world sequence-based prediction scenarios.

Beyond simple contact counts, our model accurately captures the morphology of the protein fold.

As illustrated in Figure 7, the model effectively localizes high-density contact regions, generating predictions spatially consistent with the GT. Specifically, the prediction captures the complex offdiagonal checkerboard patterns characteristic of *β*-sheet interactions and precise strand registration.

**Figure 7.**
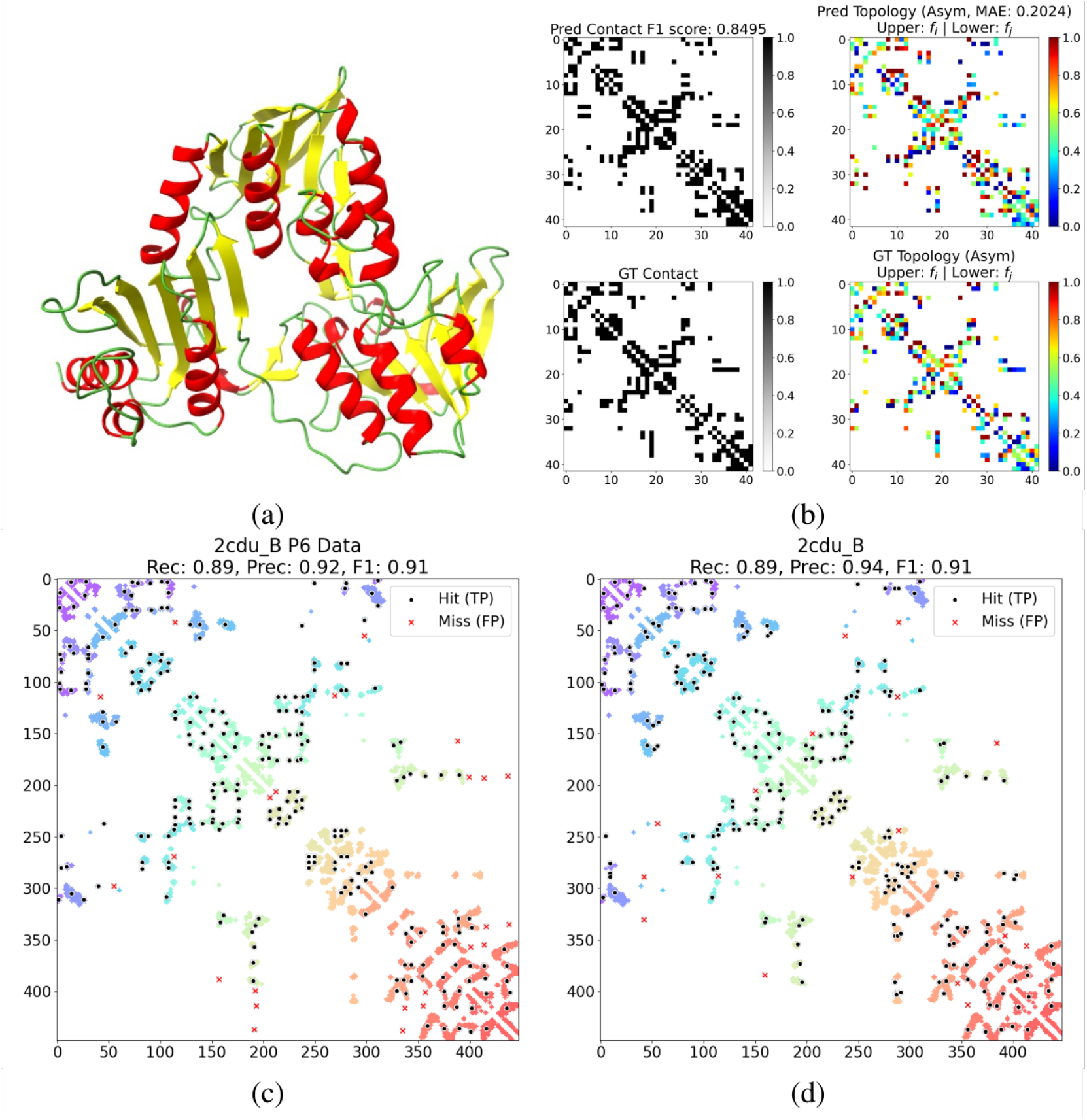
Visualization of topological prediction for protein 2cdu B. **(a)** Native 3D structural representation of the protein fold. **(b)** Detailed comparison of feature maps. The right column highlights the preservation of orientation and relative contact features (*f*_*i*_/*f*_*j*_). **(c)** Prediction based on SSEs derived from Porter 6, demonstrating robustness against input noise. **(d)** Prediction based on experimental SSEs. Black dots represent True Positives (TP); red crosses indicate False Positives (FP).

**Figure 8.**
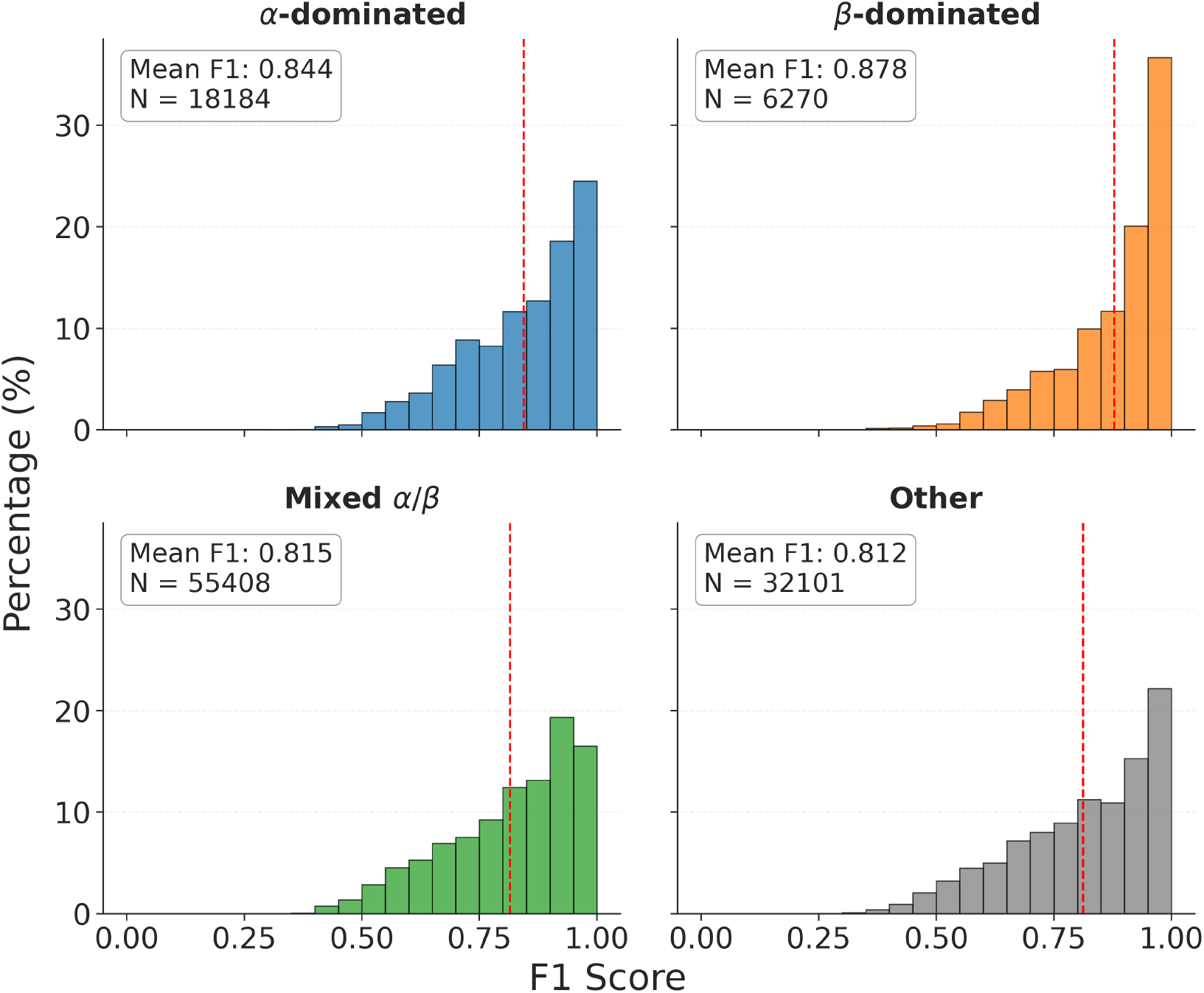
Performance distribution by secondary structure class. The histograms show the F1 scores for four distinct structural classes. The vertical red dashed line indicates the mean F1 score. Notably, *β*-dominated proteins achieve the highest accuracy (0.866), suggesting the model is effective at capturing long-range constraints.

### 2.4 Performance by Secondary Structure Class

To understand how secondary structure composition affects model performance, we classified the test proteins into four groups based on their secondary structure fractions [24]: *α*-dominated, *β*-dominated, Mix *α/β* and Others.

Our model achieved the highest performance on *β*-dominated proteins with an F1-score of 0.866. This is significantly better than the *α*-dominated class (F1-score = 0.822). This result is unexpected because *β*-sheets depend on long-range interactions, which are usually harder to predict than the local interactions found in *α*-helices. We believe this success is due to the RoPE method [25] and the attention-based framework. By capturing both absolute and relative positions, the model effectively learns relationships between distant elements [15]. Therefore, our model captures strong global constraints better than traditional models that focus on local patterns.

The performance is slightly lower for the Mixed *α/β* type, likely due to their complex topologies. Most large and multi-domain proteins belong to this category, and they often show more structural flexibility. The ‘Other’ category contains proteins with many random coils. These structures are inherently flexible and lack stable topological constraints, making them the most challenging to predict.

### 2.5 Uncertainty Quantification

Leveraging the inherently probabilistic nature of the generative flow matching model, we evaluated the correlation between predictive uncertainty, prediction accuracy, and intrinsic protein structural flexibility. For each input SSE sequence, we generated *N* = 20 independent samples to compute the pixel-wise entropy of the predicted contact map.

Given the high sparsity of contact maps, standard global metrics are often biased by the vast number of non-contact background pixels (True Negatives). To mitigate this, we restricted our evaluation strictly to **“active regions”**, defined as the union of the ground truth contacts and predicted contacts (where a contact is defined by a probability ≥ 0.6). This effectively isolates True Positives (TP), False Positives (FP), and False Negatives (FN) for analysis.

As illustrated in Figure 9(a), the model demonstrates excellent calibration. Notably, the median entropy for correctly predicted contacts (TP) is effectively zero, indicating that the model is absolutely confident in the vast majority of its accurate predictions. The elevated mean in this group is driven by a right-skewed “long tail” of challenging, ambiguous contacts. In sharp contrast, erroneous predictions (FP and FN) exhibit broadly elevated uncertainty across the entire distribution. Statistical analysis confirms a large, highly significant separation between these predictive outcomes (Mann-Whitney U test, *p <* 0.001; Cohen’s *d* = 0.876). This stark contrast verifies that high entropy serves as a highly reliable indicator of potential errors, allowing for the effective filtration of unreliable predictions.

**Figure 9.**
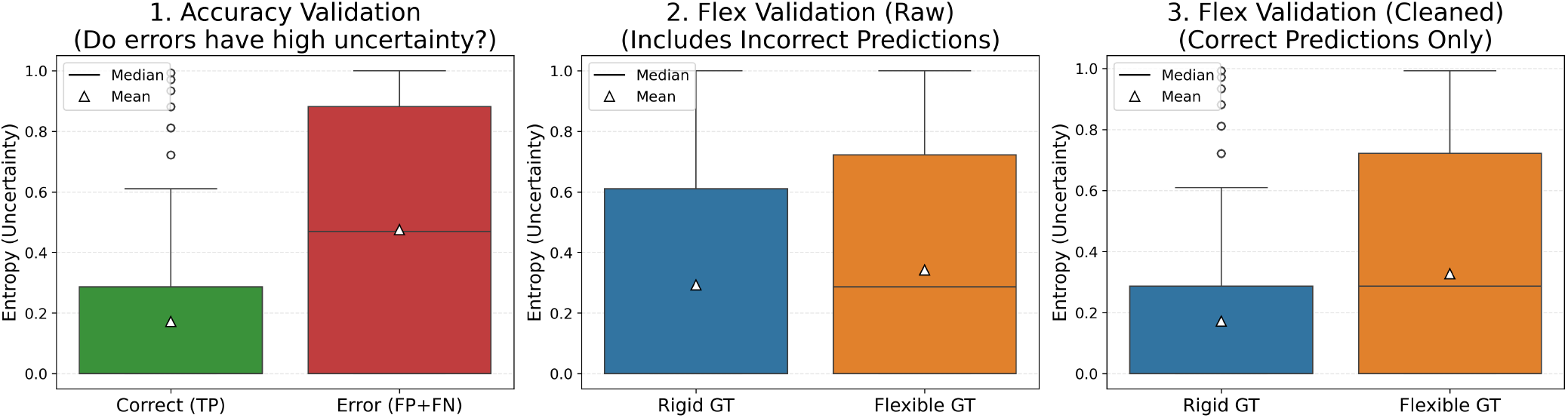
Analysis of Predictive Uncertainty regarding Accuracy and Structural Flexibility. In all boxplots, the black horizontal line represents the median, while the white triangle denotes the mean. **(a)** Uncertainty vs. Accuracy: Boxplots of predictive entropy for correct predictions (TP) versus errors (FP + FN) within active regions. The model exhibits absolute confidence (median entropy *≈* 0) for the vast majority of correct predictions, with the mean being pulled up by a long tail of uncertain outliers. **(b)** Uncertainty vs. Flexibility (All predictions): Comparison of predictive entropy between rigid (conserved) and flexible (diverse) contact regions using all predictions. The overlapping distributions indicate that prediction errors confound the distinction between rigid and flexible regions in the raw output. **(c)** Uncertainty vs. Flexibility (Correct predictions only): Entropy comparison conditioned on correctly predicted contacts. Flexible regions exhibit distinctively higher mean and median entropy than rigid regions, reflecting the model’s physical awareness of structural diversity. **Conclusion:** Overall, the model demonstrates extreme confidence in its accurate predictions and effectively differentiates intrinsically flexible regions from rigid folding cores.

Furthermore, we investigated the model’s ability to capture structural flexibility. Initially, a global comparison between rigid and flexible regions (defined by ground truth diversity) showed overlapping entropy distributions with a negligible effect size (Cohen’s *d* = 0.149, *p* = 0.0128) (Figure 9(b)), likely due to the confounding effect of high-entropy prediction errors in difficult targets. However, when we conditioned the analysis strictly on correctly predicted contacts (Figure 9(c)), a clear and statistically robust distinction emerged (Cohen’s *d* = 0.505, *p <* 0.001): flexible regions exhibit consistently higher uncertainty than rigid hydrophobic cores. This indicates that our model goes beyond static pattern matching; it successfully learns to differentiate between highly stable topological constraints and intrinsically flexible regions, thereby physically mirroring the true conformational landscape of the protein.

## 3 Discussion and Conclusion

In this work, we presented a generative framework that recovers the global topological logic of protein structures solely from compressed Secondary Structure Elements (SSEs) [11, 12]. By integrating a BERT-style attention mechanism[15] with Generative Flow Matching[14], our model transcends deterministic prediction, capturing the inherent conformational flexibility of the protein landscape.[13].

A central finding of this study is the counter-intuitive efficacy of coarse-grained SSEs in capturing long-range interactions. While traditional residue-level methods often degrade in accuracy as sequence separation increases, our model achieves its highest performance in the long-range regime and on *β*-dominated structures [24]. This challenges the reliance on atomic-level evolutionary information, suggesting that the pairwise organization of SSEs acts as a robust “topological fingerprint.” The high information density of SSEs allows the attention mechanism to bypass local noise and directly attend to global constraints, effectively solving the “folding puzzle” from the outside in [5]. The fidelity of the recovered topology extends beyond simple contact overlap. Our analysis using Circuit Topology (CT) demonstrates that the model does not merely memorize local packing patterns but learns the hierarchical logic of folding. The high recall on complex “Cross” (X) topologies—which represent the most entropically costly entanglements—indicates a learned understanding of the global energy landscape [5].

Furthermore, the mapping from coarse-grained predictions to residue-level coordinates reveals a “sub-helical precision” regime. The mean spatial alignment error of 2.69 residues falls well below the physical threshold of a single *α*-helical turn (3.7 residues) [22]. This implies that even without explicit atomic coordinates, the generative model creates a spatial probability density that locks the interaction interface with near-atomic fidelity. Finally, the probabilistic nature of Flow Matching provides a physical proxy for structural stability. We observed a distinct decoupling between generative noise and topological signal: while the model exhibits higher entropy in flexible loop regions, it maintains low-entropy, high-precision predictions for the hydrophobic core [13]. This mirrors the physical reality of proteins as thermodynamic ensembles rather than static structures. However, this capacity has limits; the model struggles to accurately distinguish flexible regions from background noise in highly disordered proteins (coils) or complex multi-domain interfaces [19]. Future work may involve calibrating the flow matching guidance scale to better balance the trade-off between generating diverse conformational ensembles and stabilizing the rigid topological core [14].

In summary, we demonstrate that the protein folding problem can be effectively reduced to a topological constraint satisfaction problem defined by SSEs. This framework offers a computationally efficient, physically interpretable alternative to end-to-end atomic prediction. Crucially, the extreme speed of this pipeline—capable of predicting 1,000 contact maps from amino acid sequences in under two minutes—enables the exploration of the protein structural genotype-phenotype (GP) map [17]. By facilitating large-scale sampling of mutant sequences to identify highly conserved folding cores, this approach bridges the gap between evolutionary sequence space and the topological principles of protein folding.

## 4 Method

### 4.1 Data Selection and Dataset Partitioning

The complete RCSB Protein Data Bank (PDB) [26] was utilized as the primary data source. Protein sequences were partitioned into individual chains, and residues with missing coordinates were excluded to ensure structural integrity. To prevent leakage and ensure the model’s generalization, we performed redundancy reduction using MMseqs2 [27]. Sequences were clustered based on a 90% sequence identity threshold using the connected component algorithm. We partitioned the resulting clusters such that approximately 80% of the data was assigned to the training set, with the remaining 20% reserved for the testing set. Importantly, all sequences belonging to the same cluster were assigned to the same partition to maintain strict separation.

### 4.2 Secondary Structure Elements (SSEs) Extraction and Representation

Secondary structure assignments were initially derived from PDB coordinates using the DSSP algorithm [28]. To facilitate a coarse-grained representation, these residue-level assignments were compressed into Secondary Structure Elements (SSEs) based on the continuity of helices (*H*) and strands (*E*). We defined a structural alphabet where helices are mapped to lowercase characters and strands to uppercase characters. To balance resolution with vocabulary size, the specific token assignment is determined by segment length using a two-tiered mapping strategy:

- **Exact Mapping:** Short segments (lengths 2–10) are mapped to unique character tokens (e.g., a 2-residue strand ‘EE’ → ‘A’, a 3-residue helix ‘HHH’ → ‘b’).
- **Compressed Mapping:** To mitigate sparsity in the vocabulary, longer segments are binned. Lengths 12–30 are mapped with a step size of 2, while segments with lengths *>* 30 are mapped with a step size of 3.

This encoding scheme effectively reduces the sequence length by approximately 13-fold [11], significantly increasing the information density for topological learning.

### 4.3 Porter 6 SSE Extraction

Porter 6 [16] leverages Protein Language Models (PLMs) for high-accuracy secondary structure prediction. By utilizing ESM-2 [29] embeddings, it captures evolutionary context without the computational overhead of Multiple Sequence Alignments (MSAs). The resulting three-state predictions were mapped to our SSE representation using the encoding scheme detailed in Method 4.2.

### 4.4 SSE Contact Map and Circuit Topology Extraction

Inter-SSE contacts were identified by computing the Euclidean distances between all atoms of pairwise SSEs, using a distance threshold of 7 °A. Unlike conventional topological definitions that rely solely on backbone atoms (e.g., *C*_*α*_ or *C*_*β*_) with looser thresholds (7–10 °A), our all-atom approach provides a more granular assessment of packing interactions. This strategy captures side-chain connectivity and accounts for the conformational flexibility inherent in ensemble predictions.

By aggregating all atomic contact pairs, we quantified the contact density, which serves as a proxy for interaction strength and facilitates the identification of flexible interfaces during loss calculation. To construct the precise circuit topology, the spatial registration of each interaction was determined by calculating the centroid of the contact interface. Specifically, we computed the mean residue index of all contacting atoms for each SSE and normalized this value to the range [0, 1], thereby defining the exact relative position of the topological connection.

### 4.5 Architecture of the Model

The proposed model employs a BERT-style encoder architecture integrated with a Continuous Normalizing Flow (CNF) framework to generate protein circuit topologies from compressed SSE sequences. The architecture consists of three primary components: a Rotary Position Embedding (RoPE) enhanced Transformer encoder, a Pair Representation Projector, and a Flow Matching Head for joint distribution modelling.

#### 4.5.1 SSE Encoder with Rotary Positional Embeddings

Given an input sequence of Secondary Structure Elements *S* = *{s*_1_, *s*_2_, …, *s*_*L*_*}*, we first embed them into latent vectors *H ∈* R^*L×d*^. To capture the crucial relative positions between SSEs, we applied Rotary Positional Embeddings (RoPE) [25] into the self-attention mechanism. Unlike absolute positional encodings [30], RoPE encodes position information by rotating the query and key vectors in the complex plane. For a token at position *m* and a feature dimension *d*, the embedding is transformed as:

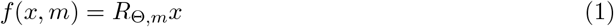

where *R*_Θ,*m*_ is a block-diagonal rotation matrix. This ensures that the attention score between token *m* and *n* depends solely on their relative distance *m* − *n*:

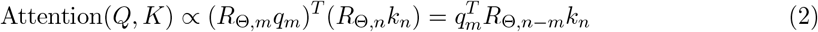

This property is particularly advantageous for topological prediction, as circuit topology is fundamentally invariant to absolute translation but sensitive to the relative arrangement of structural elements.

#### 4.5.2 Pair Representation and Joint Modeling

To predict the 2D contact map and topological features from the 1D sequence, we project the encoder representations into a pair-wise feature space, similar to the Evoformer module in AlphaFold2 [31]. The pair representation 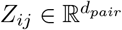 for SSEs *i* and *j* is derived via:

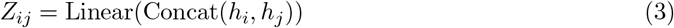

Our model unifies them into a 3-channel joint distribution *X ∈* R^3*×L×L*^. Channel 0 represents the “structural existence” (contact probability), while Channels 1 and 2 encode the asymmetric topological fractional coordinates (*f*_*i*_, *f*_*j*_). This unified representation allows the flow model to capture the correlations between contact formation and specific topological orientations.

#### 4.5.3 Generative Flow Matching Training

We adopt the Flow Matching framework [14] to model the probability density path from a standard Gaussian noise distribution *p*_0_(*x*) = *N* (0, *I*) to the data distribution *p*_1_(*x*). We define an Optimal Transport (OT) conditional vector field *u*_*t*_(*x*|*x*_1_) that pushes the noise *x*_0_ towards the ground truth topology *x*_1_ along a straight trajectory:

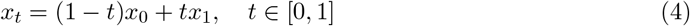

The model is trained to predict the velocity field *v*_*θ*_(*x*_*t*_, *t, c*) conditioned on the SSE sequence context *c*. The objective function minimizes the regression loss between the predicted velocity and the target drift (*x*_1_ − *x*_0_):

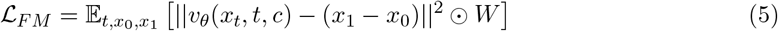

where *W* is a weighting matrix derived from contact density to balance the loss contributions from sparse contact regions and the background.

The weight of the contact matrix is determined by the number of atomic contact pairs (*N*_*ij*_) extracted as described in Method 4.4. To accommodate flexible predictions for weak interactions while enforcing consistency for rigid regions, we implemented a density-dependent weighting scheme (Eq. 6). We assigned a baseline weight of 1.0 to the background. For positive contacts, the weights were modulated using a Hill-function scaled logarithmic formulation:

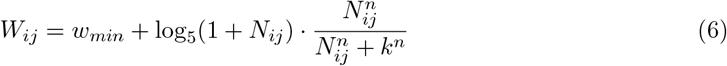

where *w*_*min*_ = 0.2 serves as the floor weight for detected contacts. The Hill coefficient *n* = 3 and half-saturation constant *k* = 50 effectively act as a soft filter: sparse contacts (*N*_*ij*_ *≪ k*) are assigned weights close to *w*_*min*_ (0.2), treating them as flexible or noisy features, whereas dense contacts (*N*_*ij*_ *≫ k*) are significantly up-weighted (*>* 1.0) to penalize deviations in the stable structural core.

During inference, we employ Classifier-Free Guidance (CFG) [32] to enhance the fidelity of the generated topology. The guided velocity field 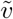 is computed as:

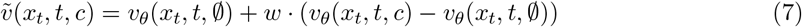

where *w >* 1 is the guidance scale. The final topology is obtained by solving the Ordinary Differential Equation (ODE) 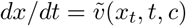 from *t* = 0 to *t* = 1 using an Euler solver.

#### 4.6 Evaluation Metrics and Analysis Protocols

To rigorously assess the model’s capability in recovering both global topological logic and local structural details, we employed a multi-level evaluation framework ranging from coarse-grained topology to residue-level coordinates.

#### 4.6.1 Circuit Topology Construction and Classification

We adopted the Circuit Topology (CT) framework [3] to classify pairwise contacts into fundamental topological arrangements. For any two contact pairs (*i, j*) and (*k, l*) (where indices represent SSE positions along the sequence), their topological relation is classified as:

- **Series (S):** Contacts are sequentially separated, *i < j < k < l*.
- **Cross (X):** Contacts are entangled, *i < k < j < l*.
- **Parallel (P):** One contact nests within another, *i < k < l < j*.

This classification transforms the 2D contact map into a topology matrix *T*, providing a robust descriptor invariant to local deformations.

#### 4.6.2 Topological Random Null Model and Sequential Constraints

To establish a baseline for interpreting the observed topological distributions, we first consider a theoretical null model based on random contact formation. For any two independent contacts defined by four distinct SSE nodes *{i, j, k, l}*, there are 4! = 24 possible linear permutations along the sequence. Based on the arrangements defined in the Circuit Topology framework:

- **Series (S):** 8 permutations (e.g., *i < j, k < l* and no overlap).
- **Parallel (P):** 8 permutations (e.g., one contact nested within another).
- **Cross (X):** 8 permutations (e.g., interleaved contacts).

In a purely uniform random system where any node can contact any other node with equal probability, the likelihood of forming each arrangement is identical: *P* (*S*) = *P* (*P*) = *P* (*X*) = 1*/*3.

However, in real protein chains, contacts are heavily constrained by sequence separation. It is well-observed that the probability of contact formation significantly decreases as the distance along the sequence increases (*P*_*contact*_ *∝ s*^−*c*^, where *s* is the separation). Under this constraint, the formation of complex entanglements like the Cross (X) topology becomes statistically rare. This is because a Cross arrangement requires two simultaneous long-range interactions to interleave their endpoints precisely. Consequently, in a constrained polymer chain, we expect the theoretical frequency of Cross topologies to be substantially lower than the 33.3% predicted by the random model (often *<* 10% depending on the chain length).

By comparing our model’s predictions against this random baseline, we can determine whether the recovered “Cross” signals represent specific, biologically relevant folding motifs or are merely artifacts of random proximity.

#### 4.6.3 Topological Similarity Metrics

We quantified the agreement between the Ground Truth (GT) and Predicted topologies using two complementary metrics:

##### Jaccard Index

To measure the overlap of topological relations, we computed the Jaccard index on the CT matrices:

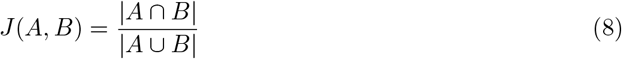

where *A* and *B* represent the sets of non-zero entries in the GT and Predicted CT matrices, respectively.

##### Damerau-Levenshtein (DL) Distance

To capture the sequential integrity of the folding path, we converted the contact maps into linear topological strings by ordering contact events along the backbone. We utilized the Damerau-Levenshtein distance [33], which accounts for insertions, deletions, substitutions, and transpositions of adjacent characters. The similarity score is normalized as:

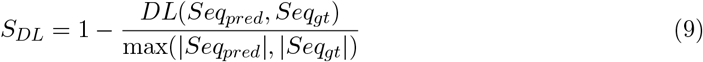

We computed this at two resolutions: *Macro-DL* (based on SSE indices) and *Micro-DL* (based on residue-level weighted centers).

#### 4.6.4 Robust Residue-Level Reconstruction

A direct pixel-to-pixel comparison between coarse-grained predictions and high-resolution ground truth can be misleading due to atomic fluctuations. To address this, we developed a robust reconstruction protocol (Figure 6 methodology) that prioritizes strong interaction cores:

##### Ground Truth Processing

We applied morphological filtering to the raw PDB contact maps. Binary dilation was performed to bridge gaps between proximal atomic contacts, followed by Connected Component Analysis (CCA) to cluster these contacts into coherent interaction patches. Small, isolated clusters (noise) were filtered out, ensuring the evaluation focuses on physically significant interfaces.

##### Prediction Smoothing

The predicted SSE contacts and their fractional coordinates (*f*_*i*_, *f*_*j*_) were mapped to residue space to generate “seed” points. To model spatial uncertainty, we applied a Gaussian kernel smoothing (*σ* = 1.5) to these seeds, generating a continuous probability density map.

##### Strict Matched Alignment Error (MAE)

For spatial precision, we defined the Strict MAE as the Euclidean distance between a correctly predicted contact’s centre (True Positive) and the nearest centroid of the matched Ground Truth cluster. This metric quantifies the localization error in residue units.

##### Global Chamfer Distance

Global Chamfer Distance (GCD) [23] is a common metric in computer vision used to measure the similarity between two sets of points. It is applied here to quantify topological fidelity. Given the set of predicted contact entries *P* and ground truth (GT) entries *G*, GCD is defined as the sum of bidirectional mean nearest-neighbor distances. The first term quantifies generative noise (precision) based on the distance from predicted points to their nearest GT entries, while the second term measures structural recall by averaging the minimum distances from GT entries to the predicted set.

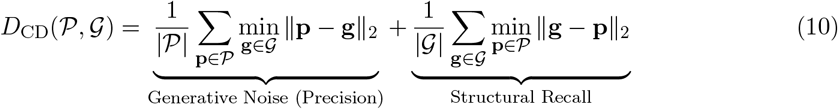

#### 4.6.5 Uncertainty Quantification

Leveraging the probabilistic nature of the Flow Matching model, we quantified predictive uncertainty by generating an ensemble of *N* = 20 samples for each input. The pixel-wise entropy *H*(*x*) was calculated as:

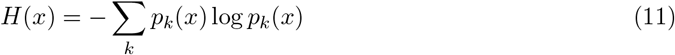

where *p*_*k*_(*x*) represents the probability of contact formation at position *x* across the ensemble. We analyzed the distribution of *H*(*x*) in rigid versus flexible regions to assess the model’s calibration.

#### 4.6.6 Secondary Structure Class Definition

Proteins were categorized based on their secondary structure composition following the thresholds defined by [24]: *α*-dominated (*α* ≥ 40%, *β* ≤ 5%), *β*-dominated (*β* ≥ 40%, *α* ≤ 5%), Mix *α/β* (both *>* 15%), and Others.

### 4.7 Code Availability

All custom code and scripts used for the SSE-based protein topology prediction are publicly available on GitHub at https://github.com/rl647/SSE_CT.

